# Structure-guided design of a targeted autoantibody degrader for neurologic disease

**DOI:** 10.64898/2026.02.01.703132

**Authors:** Marcell Zimanyi, Monica Dayao, Andoni I. Asencor, Sravani Kondapavulur, James Asaki, Krista McCutcheon, Cristina Dabaco, Aaron Bodansky, Charles S. Craik, Samuel J. Pleasure, Joseph L. DeRisi, Yifan Cheng, Michael R. Wilson, John V. Pluvinage

**Affiliations:** Department of Pharmaceutical Chemistry, University of California, San Francisco (UCSF), San Francisco, CA, USA; Chan Zuckerberg Biohub, San Francisco, CA, USA; Department of Neurology, UCSF, San Francisco, CA, USA; Weill Institute for Neurosciences, UCSF, San Francisco, CA, USA; Department of Neurological Surgery, UCSF, San Francisco, CA, USA; Biomedical Sciences Graduate Program, UCSF, San Francisco, CA, USA; Tetrad Graduate Program, UCSF, San Francisco, CA, USA; Department of Pediatrics, UCSF, San Francisco, CA, USA; Department of Biochemistry and Biophysics, UCSF, San Francisco, CA, USA; Howard Hughes Medical Institute, UCSF, San Francisco, CA, USA; Arc Institute, Palo Alto, CA, USA

## Abstract

Despite rapid progress in the diagnosis of autoantibody-mediated neurologic diseases, standard-of-care therapeutic options remain limited to nonspecific immunosuppression. Here, we report an alternative therapeutic strategy using targeted protein degradation to eliminate pathogenic autoantibodies while leaving the rest of the immune system intact. We previously discovered autoimmune vitamin B12 central deficiency (ABCD), a neurologic condition in which autoantibodies targeting the transcobalamin receptor (CD320) impair the transport of cobalamin (B12) from the blood into the central nervous system (CNS). Combining scanning alanine mutagenesis by phage display, cryo-electron microscopy, and computational modeling, we elucidated a highly conserved anti-CD320 epitope and defined the structural determinants of antigen-autoantibody binding. Next, we synthesized a lysosome-targeting chimera (LYTAC) comprising the lysosome targeting glycan, triGalNAc, fused to the antigenic epitope of CD320 as autoantibody bait. *In vitro*, this LYTAC promoted the specific lysosomal internalization and extracellular clearance of anti-CD320, restoring homeostatic cellular uptake of B12. In a passive transfer mouse model of ABCD, LYTAC treatment rapidly cleared anti-CD320 from circulation and prevented penetration of anti-CD320 into the CNS. These findings uncover the mechanism of autoantibody-antigen binding in ABCD and demonstrate targeted autoantibody degradation as a therapeutic strategy that may be generalizable to other autoimmune neurologic diseases.

## Introduction

Autoimmune neurology is a rapidly expanding field driven in part by the discovery of autoantibodies that target every level of the nervous system.^1^ For example, autoantibodies targeting the NMDA receptor on neurons can cause rapidly progressive psychosis, seizures, cognitive decline, and coma.^2^ Since the initial discovery of anti-NMDA receptor encephalitis approximately two decades ago,^3^ dozens of new autoantibody-mediated neurologic diseases have been uncovered.^4^ Despite this diagnostic revolution, therapeutic options remain limited. Nonspecific immunosuppression with corticosteroids, B-cell depletion, or plasma exchange is the current standard of care, but these treatments increase the risk of infection and are often only partially effective.^5^ Although time to diagnosis has shortened and immunosuppressive medication use has increased for several autoimmune encephalitides, patient outcomes have only marginally improved and relapse rates and mortality remain major concerns.^6^ Thus, better treatments for autoantibody-mediated neurologic diseases are needed.

Several therapies are emerging to fill this gap. Deep humoral immune system depletion and immune “reset” with CD19- and/or BCMA-targeting chimeric antigen receptor (CAR)-T cells or bispecific T-cell engagers have achieved promising results in systemic lupus erythematosus (SLE), systemic sclerosis, myositis, and stiff person syndrome.^7,8,9,10,11^ However, these therapies can cause severe and sometimes irreversible neurotoxicity, potentially limiting their utility for CNS indications.^12,13^ Rather than eliminate all B-cells, chimeric autoantibody receptor (CAAR)-T cells aim to eliminate only those expressing autoantigen-specific B-cell receptors (BCRs) and have shown promise in preclinical models of anti-NMDA receptor encephalitis and myasthenia gravis.^14,15,16^ Still, the resource-intensive process of generating a cell therapy for a rare disease and the inability of CAAR-T to eliminate circulating autoantibodies or antibody-secreting plasma cells are important translational obstacles.^17^ Fragment crystallizable (Fc) focused therapies such as neonatal Fc (FcRn) blockers and antibody-sculpting enzymes have the potential to neutralize circulating IgG, but they do not discriminate between pathogenic and protective antibodies.^18,19^ In contrast, anti-idiotype antibodies or decoy peptides target only pathogenic autoantibodies, but these occupancy-based modalities pose several pharmacologic challenges to achieve effective inhibition.^20,21^ Extracellular targeted protein degradation (eTPD) is an alternative strategy that employs event-driven pharmacology to permanently eliminate cell-surface or secreted proteins to treat disease.^22,23^ eTPD has been used to degrade checkpoint proteins from the surface of cancer cells,^24,25^ to clear total IgE from circulation in models of hypersensitivity,^26^ and even to eliminate antigen-specific pathogenic autoantibodies in models of autoimmune heart disease and myasthenia gravis.^27,28^ These versatile molecules are a promising modality for the treatment of autoantibody-mediated neurologic disease.

We recently discovered autoantibodies targeting the transcobalamin receptor (CD320) that impair the transport of vitamin B12 from the blood into the central nervous system (CNS), termed autoimmune B12 central deficiency (ABCD).^29^ These patients can develop neurologic manifestations of B12 deficiency (e.g. cognitive decline, spinal cord dysfunction) despite a normal serum B12 concentration. Anti-CD320 is surprisingly prevalent and targets a conserved region in the extracellular domain across patients with ABCD.^29,30,31^ However, the exact residues necessary for antigen-autoantibody binding remain unknown. Furthermore, although B12 supplementation is associated with partial clinical improvement, persistent circulating autoantibodies may impair homeostatic B12 transport into the CNS and limit recovery. Here, we set out to define the structural determinants of anti-CD320 binding towards the design of a targeted autoantibody degrader for ABCD.

## Results

### Convergent genetic and structural determination of an autoantibody epitope

We previously found that CD320 autoantibodies from patients with ABCD bind a conserved region in the extracellular domain of the receptor. However, the borders of this region and critical residues required for anti-CD320 binding were unknown. To map these determinants, we used three orthogonal approaches to define the anti-CD320 epitope. In the first approach, we designed a custom phage display library containing peptides tiling CD320 with a 10-amino-acid sliding window in which wild-type residues were replaced with alanine residues (Figure 1A). Using phage immunoprecipitation sequencing (PhIP-seq)^32^ with this custom mutagenesis library, we identified a 20-amino-acid region in the extracellular domain of CD320 critical for autoantibody binding in serum and cerebrospinal fluid (CSF) from 5 different patients with ABCD (Figure 1B). To increase resolution, we designed an additional phage display library comprising CD320 peptides with a single alanine mutation at every position. This revealed a 6-amino-acid epitope (SVXSLR) containing an internal threonine dispensable for anti-CD320 binding (Figure 1C). CSF antibodies from patients 4 and 5 demonstrated some degree of epitope spreading to adjacent residues, but the core epitope was conserved across all patients.

**Figure 1.**
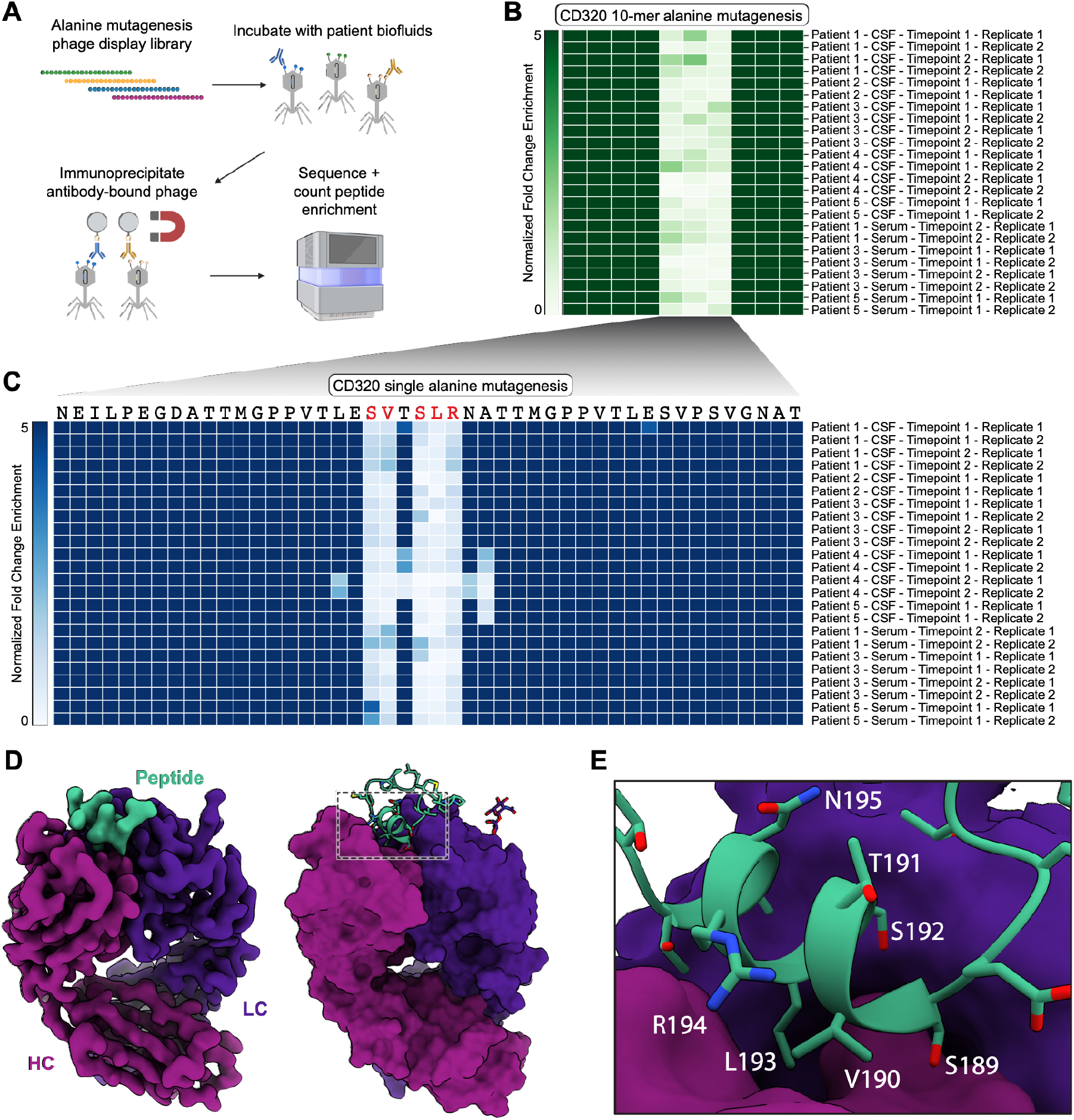
Genetic and structural determinants of autoantibody-antigen binding. **(A)** Schematic of alanine scanning mutagenesis by PhIP-seq. **(B)** Heatmap showing row-normalized fold change enrichment of peptides (columns) containing a 10 amino-acid stretch of alanine substitutions tiling across WT CD320 immunoprecipitated by antibodies in patient serum or CSF (rows). **(C)** Heatmap showing row-normalized fold change enrichment of peptides (columns) containing a single amino-acid alanine substitutions tiling across WT CD320 immunoprecipitated by antibodies in patient serum or CSF (rows). **(D)** Cryo-EM volume map of CD320 peptide (cyan) in complex with the heavy (pink) and light (purple) chains of anti-CD320 Fab. **(E)** CD320 peptide epitope (cyan) in Fab binding pocket.

To elucidate how antibodies recognize this highly conserved epitope, we determined the structure of the CD320 autoantibody-antigen complex using cryo-electron microscopy at 2.7 Å resolution (Supplementary Figure 1A; Figure 1D). The complex was formed by combining a 25-amino acid peptide comprising the PhIP-seq predicted epitope with the antigen-binding fragment (Fab) cleaved from a recombinant anti-CD320 IgG originally cloned from a CD320-specific memory B-cell from patient 1. When bound to the anti-CD320 Fab, the epitope peptide is helical, forming a short 2-turn motif spanning residues S189 to T197 (Figure 1E). Notably, residue T191 points away from the antibody interface, explaining why anti-CD320 binding is insensitive to alanine mutagenesis at this position. An N-linked glycosylation site at residue N195 is similarly oriented away from the anti-CD320 binding interface, indicating that the antibody may accommodate the glycosylated form of the protein.^33^ Taken together, these genetic and structural approaches elucidated a highly conserved epitope and suggested that antigen-specific therapies focused on this epitope might be broadly applicable across the ABCD patient population.

### Benchmarking in silico antibody epitope prediction against experimental ground truth

Recent advancements in protein structure prediction may enable rapid *in silico* determination of antibody-epitope interactions. The convergent results from our orthogonal PhIP-seq and cryo-EM experiments provided a high-confidence molecular view of this autoantibody-antigen interaction and a ground truth to which neural-network-based computational models of protein-protein interaction could be evaluated (Figure 2A).^34,35,36^ We provided AlphaFold 3 (AF3) with the amino acid sequence of the patient-derived anti-CD320 Fab along with a 15-amino-acid sliding window peptide tiling across full-length CD320 (sequence-on-sequence, Figure 2B). For each peptide, we evaluated antibody-peptide interactions using the interface predicted template modeling (iPTM) score, a confidence metric, and the estimated peptide-antibody binding energy (pyRosetta).^37^ AF3 predicted four peptides with iPTM scores above 0.8, a common threshold for high confidence interactions. Two of the four peptides contained the ground truth epitope, and these two peptides were predicted to bind anti-CD320 with the lowest binding energy (Figure 2B). We then tested this method with an experimentally determined structural starting point by providing AF3 with the empirically defined structural coordinates of the anti-CD320 Fab rather than the sequence alone (sequence-on-structure, Figure 2C). In this scenario, AF3 predicted six peptides with iPTM scores above 0.8, all of which contained the ground truth epitope (Figure 2C).

**Figure 2.**
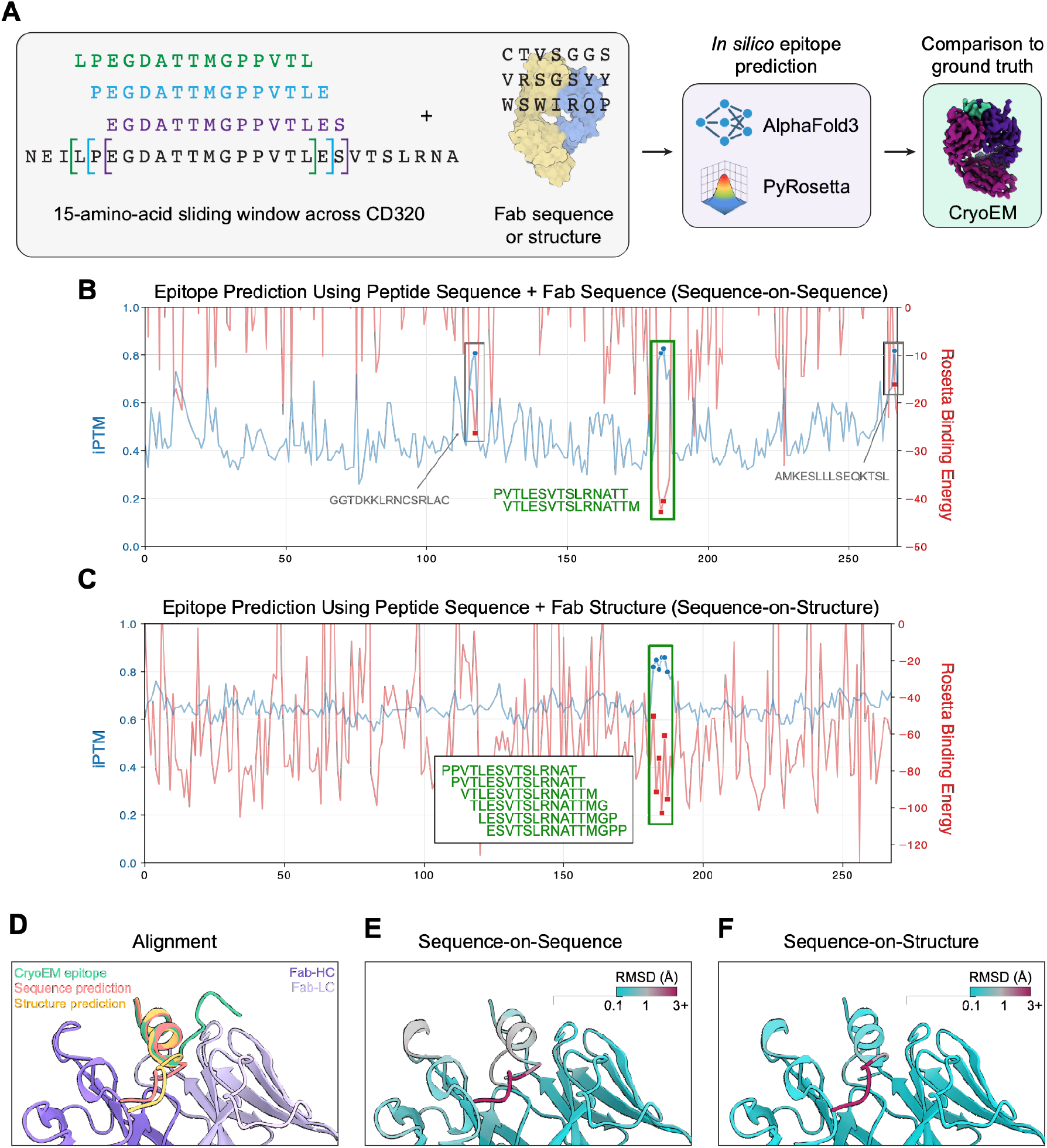
Computational modeling of autoantibody-antigen binding. **(A)** Schematic of computational modeling workflow. **(B)** AF3-predicted iPTM score (left y-axis, blue) and binding energy (right y-axis, red) of every 15-amino-acid peptide sequence tiling CD320 for the Fab sequence. **(C)** AF3-predicted iPTM score (left y-axis, blue) and binding energy (right y-axis, red) of every 15-amino-acid peptide sequence tiling CD320 for the Fab structure. **(D)** Alignment of the AF3-predicted epitopes (pink, yellow) and the cryo-EM empirical epitope (green) in complex with Fab (purple). **(E)** RMSD distance between the AF3-predicted structure with cryo-EM empirical structure for the sequence-on-sequence computational model. **(F)** RMSD distance of the AF3-predicted structure with cryo-EM empirical structure for the sequence-on-structure computational model.

In contrast, AF3 did not predict any high confidence interactions between CD320 peptides and a random Fab sequence (Supplementary Figure 2A).

Finally, we compared these AF3-predicted molecular interactions with the autoantibody-antigen cryo-EM structure via structural alignment (Figure 2D). Both the sequence-on-sequence (Figure 2E) and sequence-on-structure (Figure 2F) predictions closely aligned to our cryo-EM structure. In all 3 atomic models, the CD320 epitope peptide contained the same 2-turn helical motif with highly conserved side chain orientations. By comparing the root mean square deviation (RMSD) of Cα-carbons between each predicted structure and our determined one, we observed that the structure-on-structure prediction simulates a slight opening of the Fab complementarity determining regions (CDRs) allowing for a deeper insertion of the epitope peptide. Of note, our simulation experiments failed to predict the orientation of amino acids outside the critical binding epitope. These results suggest that both sequence-on-sequence and sequence-on-structure AF3 scanning may be practical methods for *in silico* antibody-epitope nomination followed by targeted empiric determination.

### Design of a targeted autoantibody degrader for ABCD

Several technologies have emerged for targeted degradation of extracellular proteins. These include lysosome-targeting chimeras (LYTACs) which are bifunctional molecules comprising a protein-binding arm fused to a synthetic glycan ligand for receptor-mediated endocytosis.^38^ Thus, extracellular targets are internalized and shuttled to the lysosome for degradation. Most LYTACs target proteins using antibodies. We set out to target autoantibodies using proteins as bait. Specifically, we conjugated a peptide containing the conserved antigenic epitope of CD320 to triGalNAc, a high-affinity glycan ligand of the liver-specific, lysosome-targeting asialoglycoprotein receptor (ASGPR). Our two-step bioorthogonal synthesis involved 1) strain-promoted azide-alkyne cycloaddition of three azide-modified peptides to a triantennary heterobifunctional PEG-linker, and 2) NHS ester coupling of the product of step 1 to triGalNAc with an alkyl-C5-linker (Figure 3A). We confirmed binding to both anti-CD320 and ASGPR by biolayer interferometry (Supplementary Figures 3A, B).

**Figure 3.**
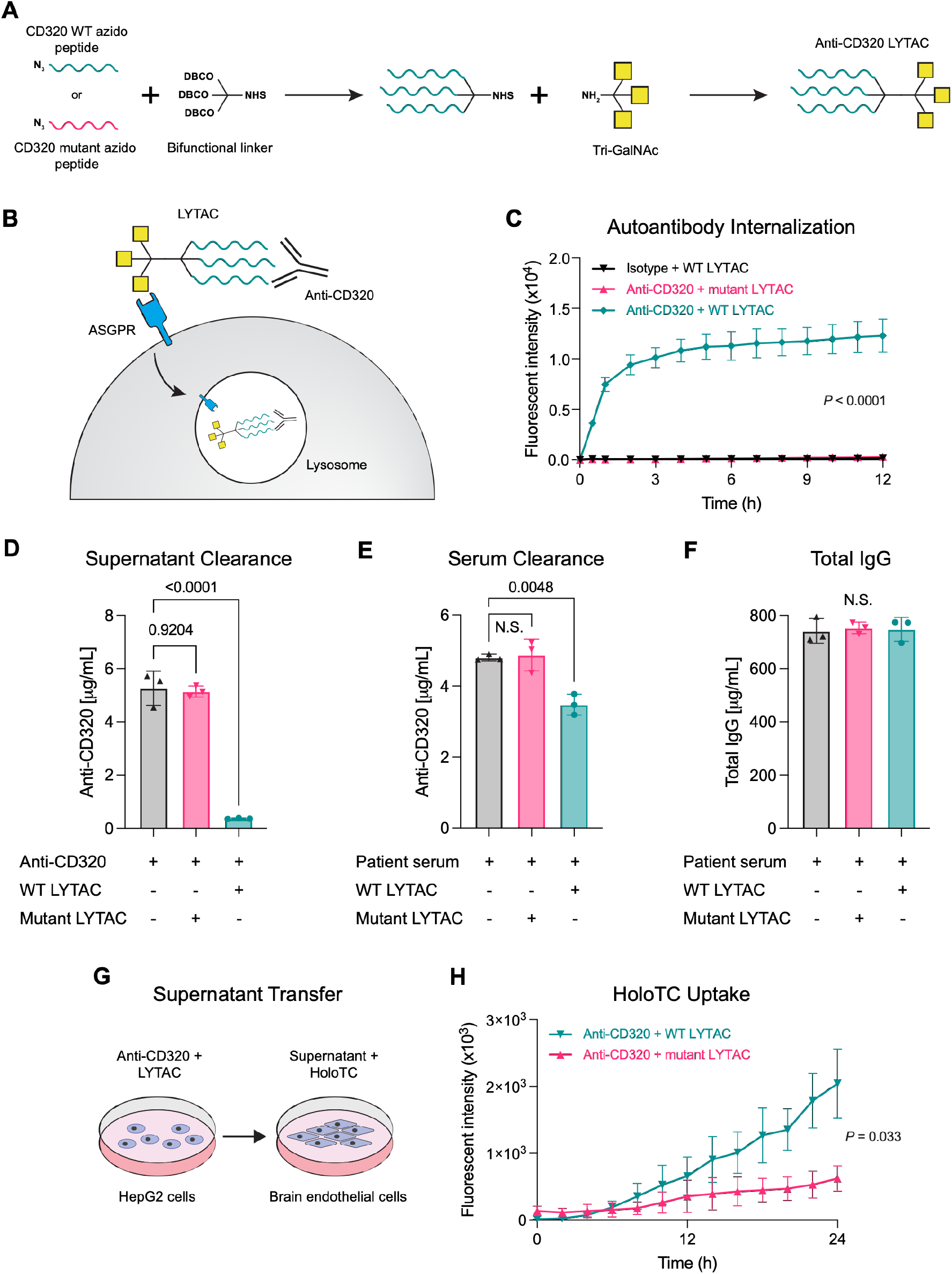
Synthesis and *in vitro* efficacy of an anti-CD320 targeting LYTAC. **(A)** Synthesis of WT (teal) and mutant (pink) LYTACs. **(B)** Model of LYTAC mechanism of action. **(C)** Antibody internalization of isotype + WT LYTAC (black), anti-CD320 + mutant LYTAC (pink), and anti-CD320 + WT LYTAC (teal) in HepG2 cells measured by time-lapse fluorescence microscopy (one-way ANOVA with Tukey’s multiple hypothesis correction). **(D)** Anti-CD320 concentration in the supernatant of HepG2 cells treated with anti-CD320 alone (black), anti-CD320 + mutant LYTAC (pink), or anti-CD320 + WT LYTAC (teal) measured by Luminex assay (one-way ANOVA with Tukey’s multiple hypothesis correction). **(E)** Anti-CD320 concentration in the supernatant of HepG2 cells treated with seropositive patient serum (black), serum + mutant LYTAC (pink), or serum + WT LYTAC (teal) measured by Luminex assay (one-way ANOVA with Tukey’s multiple hypothesis correction). **(F)** Total IgG concentration in the supernatant of HepG2 cells treated with seropositive patient serum (black), serum + mutant LYTAC (pink), or serum + WT LYTAC (teal) measured by ELISA (one-way ANOVA with Tukey’s multiple hypothesis correction). **(G)** Schematic of supernatant transfer experiment. **(H)** HoloTC uptake in brain endothelial cells incubated with the supernatant of HepG2 cells treated with anti-CD320 + mutant LYTAC (pink) or anti-CD320 + WT LYTAC (teal) measured by time-lapse fluorescence microscopy (two-way t-test).

With this LYTAC in hand, we evaluated its ability to target anti-CD320 for lysosomal degradation via ASGPR (Figure 3B). CD320-LYTAC promoted rapid internalization and lysosomal trafficking of anti-CD320 into a human liver cell line (HepG2) (Figure 3C). In contrast, CD320-LYTAC had no effect on the internalization of an isotype control antibody, and a LYTAC containing a mutant CD320 peptide (mutant-LYTAC) had no effect on the internalization of anti-CD320, confirming target specificity. To orthogonally validate this finding, we measured anti-CD320 in the supernatant of LYTAC-treated HepG2 cells using an epitope-specific immunoassay. After 24 hours, anti-CD320 was almost completely cleared from the supernatant of cells treated with the WT-LYTAC but remained present at high concentrations in the supernatant of mutant-LYTAC treated cells (Figure 3D). Dose-response analysis demonstrated that WT-LYTAC cleared the upper range of physiologic anti-CD320 concentration with an IC50 of ∼47 nanomolar (Supplementary Figure 3C).

Pathogenic autoantibodies exist as a minor constituent of complex polyclonal immunoglobulin repertoires in the serum. Many of these immunoglobulins protect against infections. Thus, targeted elimination of pathogenic autoantibodies while sparing protective bystanders may effectively treat autoimmune disease without subjecting patients to the risks of immunosuppression. To evaluate the specificity of CD320-LYTAC in vitro, we treated HepG2 cells with serum from a patient with ABCD. After 24 hours, CD320 LYTAC reduced anti-CD320 concentration by ∼27% (Figure 3E) without depleting total IgG concentration (Figure 3F). This reduction was less pronounced than LYTAC-mediated clearance of recombinant anti-CD320, possibly due to competition with endogenous ASGPR ligands in human serum.^39^

Anti-CD320 is hypothesized to impair the cellular uptake and transport of B12 at the blood-brain barrier. To evaluate whether LYTACs can reverse this phenotype *in vitro*, we first treated HepG2 cells with anti-CD320 and WT- or mutant-LYTACs (Figure 3G). Next, we transferred the supernatant from the first culture onto primary human brain endothelial cells and spiked in the bioactive form of B12 bound to its carrier protein, holotranscobalamin (holoTC). HoloTC was labeled with a pH-sensitive fluorescent dye that emits a red signal when internalized and trafficked to the acidic lysosome. Using time lapse fluorescent microscopy, we found that WT-LYTAC treatment restored cellular uptake of holoTC *in vitro* (Figure 3H). These results suggest that targeted anti-CD320 degradation can reverse the cellular pathophysiology of ABCD.

### Targeted autoantibody degradation in a passive transfer model of ABCD

In many autoimmune neurologic diseases, the pathogenic autoantibody is present at higher concentrations in the serum than in the CSF.^40^ While intrathecal antibody synthesis has been observed in several of these conditions,^41^ others, including anti-AQP4 in neuromyelitis optica spectrum disorder (NMOSD),^42^ anti-MOG in myelin oligodendrocyte glycoprotein-associated disease (MOGAD),^43^ and anti-CD320 in ABCD, are thought to derive from antibody-secreting cells in the periphery. Indeed, plasma exchange is therapeutic in these conditions, suggesting that clearance of pathogenic autoantibodies in the periphery is sufficient to abrogate disease activity in the CNS.

To evaluate the therapeutic potential of CD320-LYTAC *in vivo*, we developed a passive transfer mouse model of ABCD (Figure 4A). Due to the limited cross-reactivity of human patient-derived anti-CD320 with mouse CD320, this model was not designed to recapitulate the phenotype of ABCD. Rather, we sought to evaluate autoantibody clearance and tissue distribution. Mice were treated anti-CD320 and LYTAC or PBS via intraperitoneal injection. The autoantibody was labeled with a fluorescent dye to enable tracking of tissue penetrance. Circulating levels of anti-CD320 were assessed by serial blood collection, followed by terminal organ collection at day 4. By 24 hours, LYTAC treatment rapidly cleared circulating anti-CD320 approximately 10-fold compared to PBS treated control mice (Figure 4B). By 72 hours, circulating anti-CD320 was undetectable in all LYTAC-treated mice but sustained at a high concentration in control mice. Using fluorescent microscopy, we found that LYTAC treatment reduced penetrance of anti-CD320 into the brain parenchyma (Figure 4C, Figure 4D). These findings demonstrate proof-of-principle efficacy and suggest that CD320 LYTAC may be a viable therapeutic strategy to eliminate pathogenic autoantibodies in ABCD.

**Figure 4.**
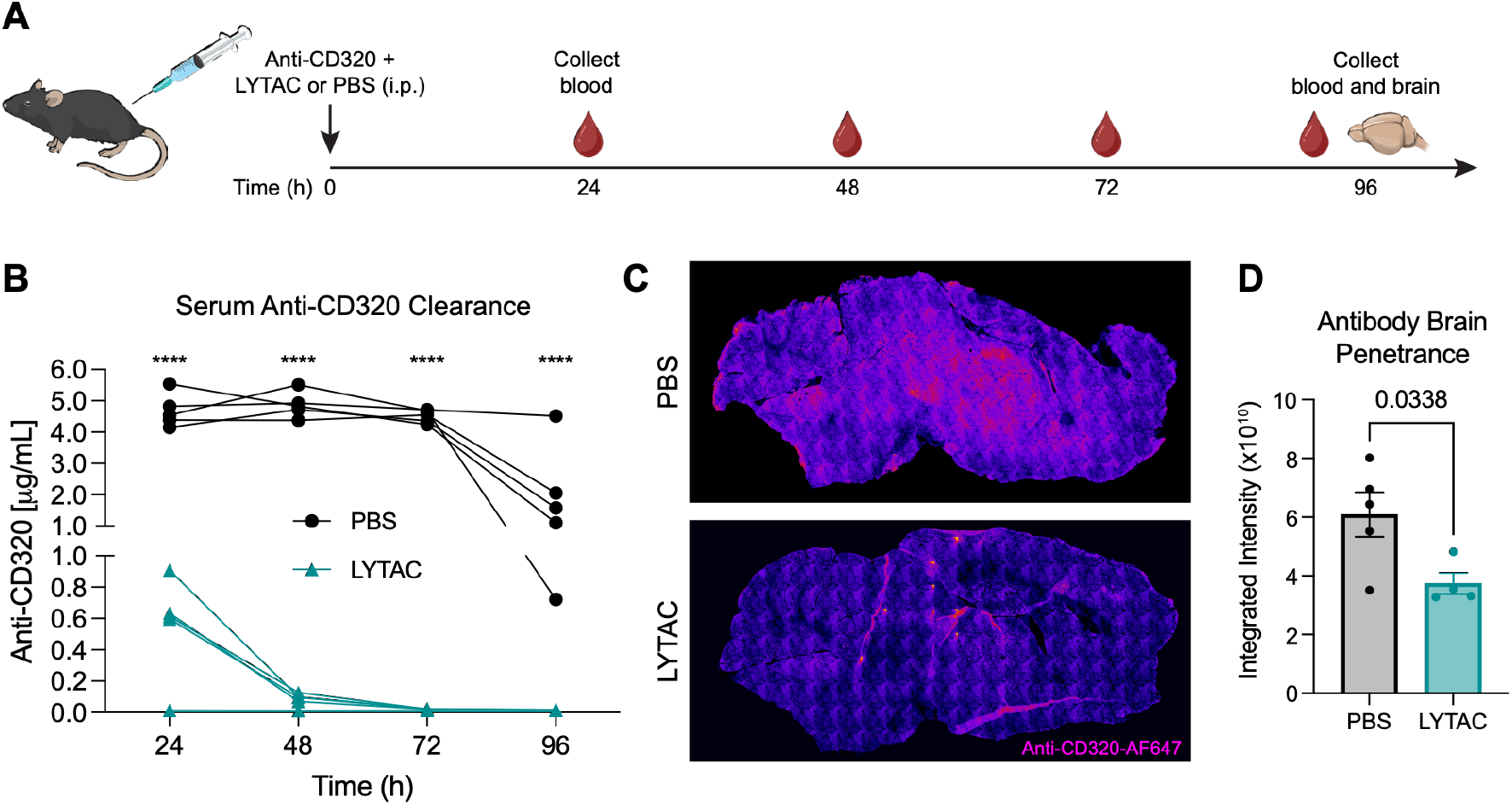
*In vivo* evaluation of an anti-CD320 targeting LYTAC. **(A)** Schematic of in vivo autoantibody passive transfer experiment. **(B)** Anti-CD320 concentration in the serum of mice treated with anti-CD320 LYTAC (teal) or PBS (black) measured by Luminex assay (two-way ANOVA with Sidak multiple hypothesis correction). **(C)** Representative fluorescent images of AF647-labeled anti-CD320 (magenta, LUT) brain tissue distribution in a mouse treated with PBS (top) or LYTAC (bottom). (**D**) Total integrated fluorescence intensity of AF647-labeled anti-CD320 in PBS (black) or LYTAC (teal) treated mice (two-sided t-test +/- s.e.m.).

## Discussion

A deep molecular understanding of a disease target can inform drug development.^44,45,46,47^ Here, genetic, biochemical, and computational approaches converged to reveal the structural determinants of autoantibody-antigen binding in ABCD. This molecular view of a highly conserved autoantibody epitope informed the design of a targeted autoantibody degrader that cleared anti-CD320 while leaving total IgG concentration intact. These findings elucidate the mechanism of an emerging autoimmune neurologic disease and demonstrate a new therapeutic strategy that may be generalizable to other autoimmune neurologic conditions.

A major bottleneck in autoantibody discovery is the prospective isolation of antigen specific monoclonal antibodies from patient-derived B-cells or antibody-secreting cells.^48,49,50,51^ Using the cryo-EM structure of autoantibody-antigen as ground truth, we found that AF3 accurately nominated this interaction with sequence information alone. Although we only validated this approach for anti-CD320, it may be generalizable to other autoantibody-antigen targets for *in silico* monoclonal isolation followed by empirical biochemical validation.

Compared to the single infusion of CAR-T cells used in several immune reset clinical trials, LYTACs would likely require repeated dosing for durable and sustained autoantibody clearance. Modifications including pH-sensitive binding arms and FcRn-mediated recycling domains have recently been shown to improve the pharmacokinetic profile of LYTACs and enable their catalytic degradation potential.^52^ Thus, repeated but infrequent dosing may be possible with this emerging therapeutic modality.

In addition to durability, the safety of targeted autoantibody degradation with LYTACs will need to be evaluated in future studies. Introduction of the immunogenic epitope of CD320 on a LYTAC could theoretically exacerbate autoantibody production. Using an alternative, ubiquitously expressed lysosome-targeting receptor such as M6PR may enable the clearance of both circulating antibody as well as membrane-bound antibody (BCR) on the surface of antigen-specific B-cells.^53^ This may mitigate the possibility of stimulating antigen-specific memory B-cell clonal expansion.

This study has several limitations. First, we resolved a high-resolution structure of anti-CD320 bound to its peptide epitope, but its structure bound to full-length CD320 may reveal additional allosteric modifiers of autoantibody binding. Second, although our phage display data suggest that the anti-CD320 epitope is highly conserved across all queried patients with ABCD, we resolved the cryo-EM structure of an autoantibody derived from a single patient. Future studies with additional patient-derived anti-CD320 clones may reveal sequence variation or post-translational modifications that alter the determinants of autoantibody binding. Third, CD320 divergence between mice and humans precluded our ability to test our LYTAC in a model that recapitulates the clinical phenotype of ABCD. Humanized mice or non-human primates may provide a more clinically relevant model for therapeutic development.

Complications of treatment with nonspecific immunosuppression are sometimes as devastating as the disease itself. We demonstrate targeted autoantibody degradation as one alternative therapeutic strategy that may reduce morbidity and improve outcomes in patients with autoimmune neurologic diseases like ABCD.

## Acknowledgments

We thank Glenn Gilbert and the UCSF EM Core which is partially supported by NIH grants S10OD020054, S10OD021741, S10OD026881, and the Howard Hughes Medical Institute. This work was supported by NINDS (RM1NS138808 and R01MH122471 to S.J.P., M.R.W., J.L.D.; UE5NS070680 to J.V.P.; K08NS142575 to J.V.P.), the Westridge Foundation (to M.R.W.), the Burroughs Wellcome Fund (to J.V.P.), and Arc Institute (to J.V.P.). We thank the patients and their families for participation in this study.

## Methods

### Human sample collection

Patients were enrolled in a research study for suspected neuroinflammatory diseases (UCSF IRB# 13-12236). Serum and CSF samples were prepared neat or diluted 1:1 in antibody storage buffer (final concentration 20% glycerol, 20mM HEPES, 0.02% sodium azide in PBS) and stored at -80C.

### Scanning mutagenesis by phage display

We adapted a previously published protocol for phage immunoprecipitation sequencing (PhIP-Seq).^54,55,56^ A custom library containing 1) 49–amino acid WT peptides with 25–amino acid overlaps tiling CD320, 2) 49–amino acid peptides with 10-mer alanine substitutions tiling CD320, and 3) 49–amino acid peptides with single alanine substitutions tiling CD320 was cloned into T7 bacteriophage. Patient serum or CSF was incubated with 10^10^ plaque-forming units of the phage library, antibodies were immunoprecipitated with protein A/G magnetic beads, and antibody-bound phage was amplified in *Escherichia coli* before a second round of immunoprecipitation. Enriched phage lysates were adaptor-ligated and barcoded before pair-end sequencing on an Illumina NovaSeq to a depth of 2 million reads per sample. Reads were trimmed, aligned at the amino acid level using RAPSearch, and normalized to sequencing depth to generate reads per 100,000 (RPK) for each sample. Enriched peptides were identified by calculating the fold change (FC) of normalized counts between samples immunoprecipitated with serum or CSF versus magnetic beads only.

### Cryo-EM Sample Preparation and Data Collection

CD320 autoantibody Fab was generated by processing CD320 autoantibody IgG with a papain-based Fab preparation kit (Pierce, 44985). CD320 Fab was concentrated to 1 mg/mL and combined with CD320 tripeptide (3x MGPPVTLESVTSLRNATTMGPPVTL-NHS-PEG5-tris-PEG3-DBCO) at a 1-to-1 equimolar ratio and incubated at room temperature for 30 minutes. 3 microliters of the complex was pipetted onto glow-discharged gold grids coated in holey gold film (UltrAUfoil, 300 mesh 1.2/1.3). Grids were blotted and plunge-frozen using a Vitrobot Mark IV equipped with Whatman type 4 blotting paper with a blotting time of 4 seconds and a blot force of -2 at 4 °C and 100% humidity. Data was collected on a Thermo Fisher Krios at the UCSF Cryo-EM Center for Structural Biology, operated at an acceleration voltage of 300 kV, equipped with an XFEG, a BioQuantum energy filter (slit width set to 10 eV) and a Gatan K3 camera. A nominal magnification of 135,000x was used for a physical pixel size of 0.818 Å (0.409 in super resolution) with a total dose of 48 e-/Å^2^. Automated data collection was performed using SerialEM to collect movies with a defocus range between 0.8-2 Å.

### Cryo-EM Image Processing and model building

Dose-fractionated images stacks were motion corrected using both cryoSPARC and MotionCor2.^57,58^ CTF estimation was performed using cryoSPARC, followed by micrograph curation, blob-based particle picking, 2D classification, and ab initio modeling. The ab initio model was used to generate templates for further particle picking and curation. Iterative rounds of multi-class ab initio modeling and heterogeneous refinement were used to generate particle stacks where the Fab was clearly resolved. The final model was generated using reference-based motion correction followed by non-uniform 3D refinement. A model of the CD320 autoantibody Fab was generated by AF3 and was docked into cryo-EM density and refined using ISOLDE and PHENIX.^35,59,60^ The peptide was modeled *de novo* into the remaining cryo-EM density using coot and further refined using ISOLDE and PHENIX.^61^ Final collection, refinement, and validation statistics are reported in Supplementary File 1. Data has been deposited in the Protein Data Bank (PDB ID 9ZYU).

### AlphaFold Computational modeling of antibody-antigen interactions

To predict structures of antibody:peptide complexes, we provided AF3 with the amino acid sequence of the patient-derived anti-CD320 Fab (heavy and light chain) along with a 15-amino-acid sliding window peptide tiling across full-length CD320 (268 15-mer peptides). For each run, AF3 outputs an interface predicted template modeling (iPTM) score for the peptide (measuring the average iPTM between the peptide and the two antibody chains), measuring the model’s confidence in the predicted relative positions of the peptide to the antibody chains. Values higher than 0.8 represent high-quality predictions, values below 0.6 suggest a failed prediction, and values between 0.6 and 0.8 is considered a grey zone where predictions could be correct or incorrect (https://www.ebi.ac.uk/training/online/courses/alphafold/). We performed the same experiment with a previously published unrelated Fab (Fab3).^62^ In the experiment where the empirically derived structure of the autoantibody is given to AF3, we input mmCIF files of both the heavy and light chain into the model and set ‘unpairedMsa’ and ‘pairedMsa’ fields to empty strings (““), which is equivalent to running AF3 without the multiple sequence alignment (MSA) step. Predicted antibody:peptide structures from AF3 were used to compute interface binding energies (ΔΔG) between peptide and antibody. We used PyRosetta’s InterfaceAnalyzerMover (IAM) with the ref2015 scoring function.^37^ For three-chain peptide-antibody complexes, the peptide (chain A) was designated as one partner and the antibody heavy and light chains (chains B and C) as the other partner using the docking partners string “A_BC”. The IAM protocol calculated ΔΔG by comparing the energy of the bound complex to the sum of energies of the separated partners. To obtain binding energies that account for side-chain optimization, both the bound complex state and the separated partner states were repacked prior to energy evaluation (pack_input=True, pack_separated=True). This approach allows side chains to adopt favorable rotamers in both bound and unbound states, providing a more accurate estimate of the binding free energy change. The final ΔΔG values were reported in Rosetta Energy Units (REU), with more negative values indicating stronger binding interactions.

### LYTAC synthesis

CD320 peptides encompassing the wild-type autoantibody epitope sequence (MGPPVTLESVTSLRNATTMGPPVTL) or mut sequence (MGPPVTLEAAAAAANATTMGPPVTL) were synthesized with an N-terminal azide-modified lysine (GenScript). Each peptide was resuspended in DMSO and incubated at 4 molar excess with the heterotetrafunctional linker, NHS-PEG5-tris-PEG3-DBCO (Conju-Probe, CP-2235) for 24 hours at room temperature. Upon completion of the SPAAC reaction, the product was purified using a Zeba size exclusion column (7 kDa MWCO, ThermoFisher, 89882). Purified product was confirmed by SDS-PAGE, then incubated with an equimolar ratio of tri-GalNAc-C5-amine (Tocris, 7781) at room temperature for 2 hours in 0.1M sodium bicarbonate in PBS. Upon completion of the NHS ester coupling, the reaction was quenched with Tris and again purified using a Zeba size exclusion column (7 kDa MWCO, ThermoFisher, 89882). Purified product was confirmed by SDS-PAGE.

### Cell culture

HepG2 cells were purchased from ATCC (HB-8065) and cultured in DMEM + GlutaMAX with 10% FBS. HEK293T cells were cultured in DMEM + GlutaMAX with 10% FBS. Primary human brain microvascular endothelial cells (Cell Systems, ACBRI 376) were cultured in Complete Classic Medium (4Z0-500).

### Antibody internalization assay

Recombinant patient-derived anti-CD320 or human IgG1 isotype control antibody (BioXCell, BP 0287) were conjugated to pHrodo Red iFL STP ester (ThermoFisher, P36011) at a dye:protein molar ratio of 10:1. Free dye was removed using a 40 kDa MWCO Zeba Spin Desalting Column (ThermoFisher, A57759). HepG2 cells were lifted using TrypLE Express (ThermoFisher, 12605010) and plated on TC-treated 96 well plates at 10,000 cells per well the day prior to the assay. Labeled antibodies were diluted to 1 microgram/mL in serum free DMEM and pre-incubated with WT- or mutant CD320 LYTAC (3 micromolar) for 15 minutes at room temperature. Complete medium was removed from HepG2 cell containing wells replaced with pre-incubated medium containing antibodies and LYTACs. Antibody internalization was monitored every 1-2 hours for 12-24 hours using a live cell time-lapse fluorescent microscope (Incucyte SX5).

### Luminex immunoassay

We adapted a previously published protocol for autoantibody detection using the Luminex platform.^63^ Briefly, a peptide comprising the epitope for anti-CD320 was synthesized and conjugated to BSA via carbodiimide coupling to a C-terminal cysteine. Spectrally-distinct Luminex magnetic beads were conjugated to CD320 peptide-BSA or BSA alone via NHS coupling to free amines following the manufacturer’s protocol (Luminex, 40-50016). Serum or supernatant was incubated with Luminex beads at a final dilution of 1:500 in PBS + 0.05% Tween 20 (PBST) containing 2% non-fat milk for 1 hour at room temperature with agitation. Samples were washed two times with PBST then stained with PE-conjugated anti-human IgG Fc secondary antibody (1:2,000, BioLegend, 637310) for 30 minutes at room temperature with agitation. Beads were washed three times with PBST and analyzed on a Luminex LX200 cytometer. Quantitative accuracy of this assay was confirmed using a standard curve with a recombinant patient-derived monoclonal anti-CD320 autoantibody. Anti-CD320 concentration was interpolated from the standard curve using an asymmetric sigmoidal 5PL nonlinear regression (GraphPad).

### Biolayer interferometry

Anti-Human Fc biosensors (Gator) were loaded with 50nM of anti-CD320 or 100nM of ASGPR-Fc fusion protein (R&D Systems, 10255-AS-050) in Q Buffer (Gator) on a Gator Prime BLI instrument. Loading was terminated in the linear phase once a response of at least 1 nm was achieved. Loaded probes were then equilibrated in Q buffer before association/dissociation with 10nM of LYTAC. Curves were normalized to reference wells, aligned to baseline, and processed using Savitzky-Golay filtering.

### HoloTC internalization assay

Recombinant human transcobalamin 2 (R&D Systems, 7895-TC-050) was conjugated to pHrodo Red iFL STP ester at a dye:protein molar ratio of 4:1. Free dye was removed using a 40 kDa MWCO Zeba spin desalting column. pHrodo-conjugated transcobalamin was incubated with 3X molar excess of cyanocobalamin for 1 hour at room temperature to form holotranscobalamin. Antibody internalization was performed as described above except with unlabeled anti-CD320 or isotype control antibody. The supernatant was collected after 24h incubation on HepG2 cells, LYTAC was removed using a 40 kDa MWCO Zeba spin desalting column equilibrated in DMEM, and purified supernatant was transferred to BEC cells plated at 20,000 cells per well in a 96 well plate. After 30 minutes of pre-incubation at 37 C, 100 nanograms of pHrodo holotranscobalamin was added to each well. Holotranscobalamin internalization was monitored every 1-2 hours for 12-24 hours using a live cell time-lapse fluorescent microscope (Incucyte SX5). The average total integrated intensity across three technical duplicates was used to compare uptake dynamics between conditions.

### In vivo autoantibody passive transfer

Recombinant patient-derived anti-CD320 was conjugated to AlexaFluor 647 NHS Ester (ThermoFisher, A27573) at a dye:protein ratio of 10:1. Free dye was removed with a 40 kDa MWCO Zeba spin desalting column. Conjugated antibody was mixed with LYTAC or PBS. 200 micrograms of antibody (1mg/mL) and 20uL of LYTAC (34 micromolar) or PBS were injected into 6-month-old C57Bl/6 mice (Jax) via intraperitoneal injection. Every 24 hours for 4 days, 5 microliters of blood were collected by tail nick and mixed with 5uL of PBS containing EDTA (final concentration 5mM). Blood was centrifuged at 1000g for 2 minutes, and the acellular plasma fraction was collected and stored at -80 C. Anti-CD320 concentration was measured using the aforementioned Luminex immunoassay. Four days after injection, mice were euthanized, perfused with 4% PFA, and brains were dissected and post-fixed in 4% PFA at 4C overnight. Next, fixed tissue was cryopreserved in 30% sucrose at 4C.

### Immunofluorescence microscopy

Fixed and cryopreserved tissue was frozen in OCT, and 20 micron sections were prepared on a CryoStat and mounted onto SuperFrost slides. Slides were dried, rehydrated, stained with DAPI, and mounted using ProLong Gold antifade reagent. AlexaFluor647 signal from labeled anti-CD320 was imaged on a slide scanner (Nikon Axios) and a laser-scanning confocal microscope (Zeiss LSM780). The AF647 channel was converted to an LUT mask to visualize distribution and intensity of labeled anti-CD320 throughout the brain parenchyma.

---

**Supplementary Figure 1.**
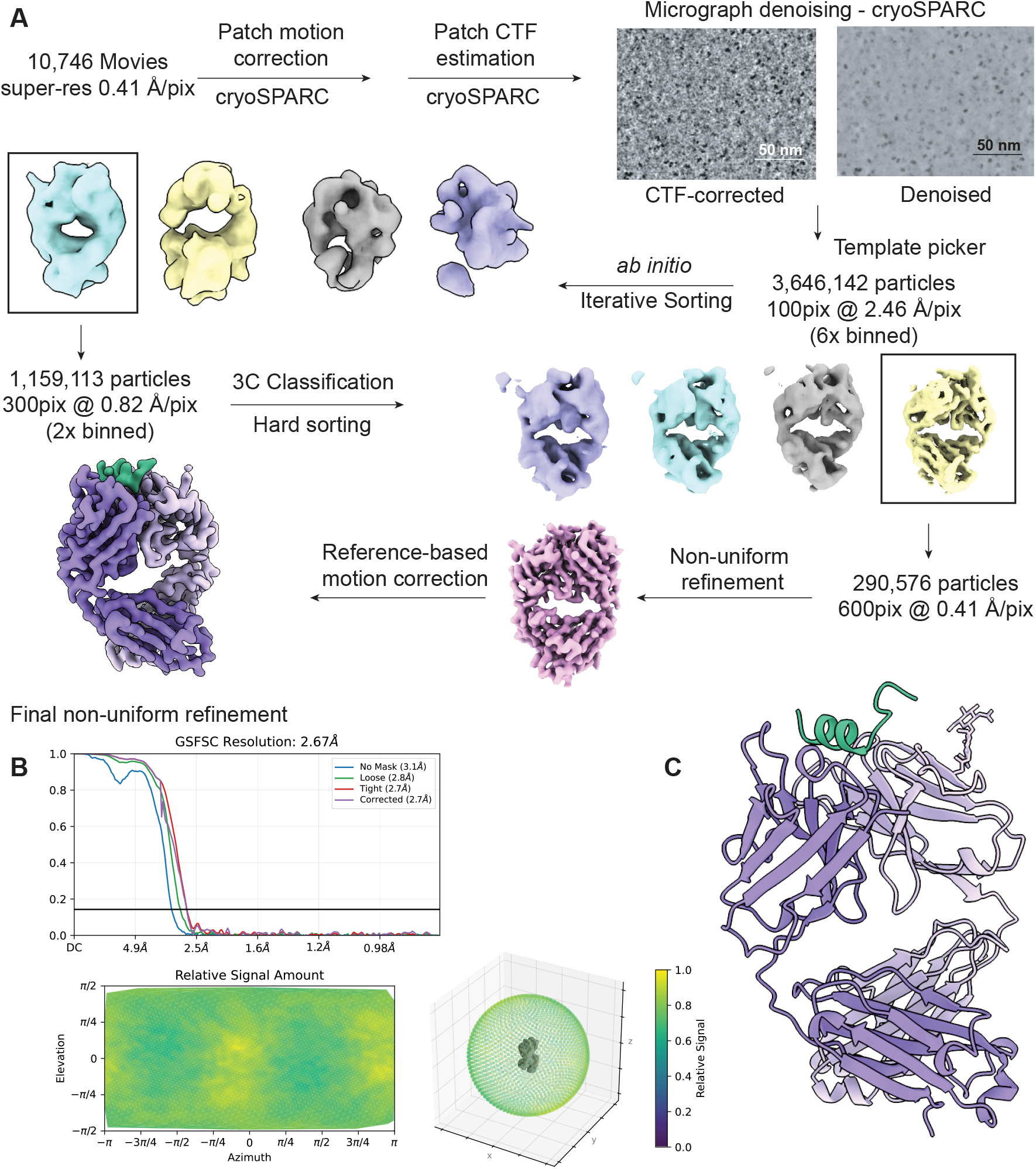
Cryo-EM workflow of anti-CD320 Fab bound to CD320 epitope peptide, related to Figure 1. (**A**) Workflow of cryo-EM data processing, including representative CTF-corrected dose-weighted micrographs before and after denoising. (**B**) Gold-standard Fourier shell correlation (FSC) curves and relative viewing angle distributions for the final refined map. (**C**) Model of an anti-CD320 Fab fragment (Light purple -- light chain, dark purple -- heavy chain) bound to the CD320 epitope peptide (green).

**Supplementary Figure 2.**
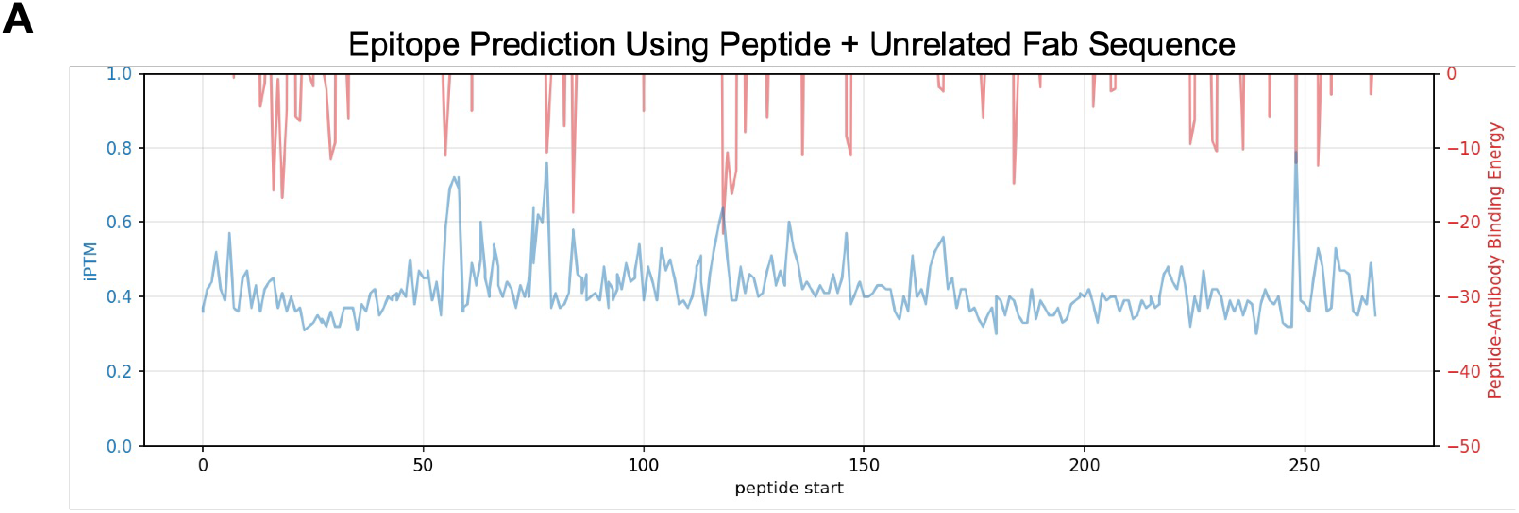
(**A**) AF3-predicted iPTM score (left y-axis, blue) and binding energy (right y-axis, red) of every 15-amino-acid peptide sequence tiling CD320 in complex with an unrelated Fab sequence.

**Supplementary Figure 3.**
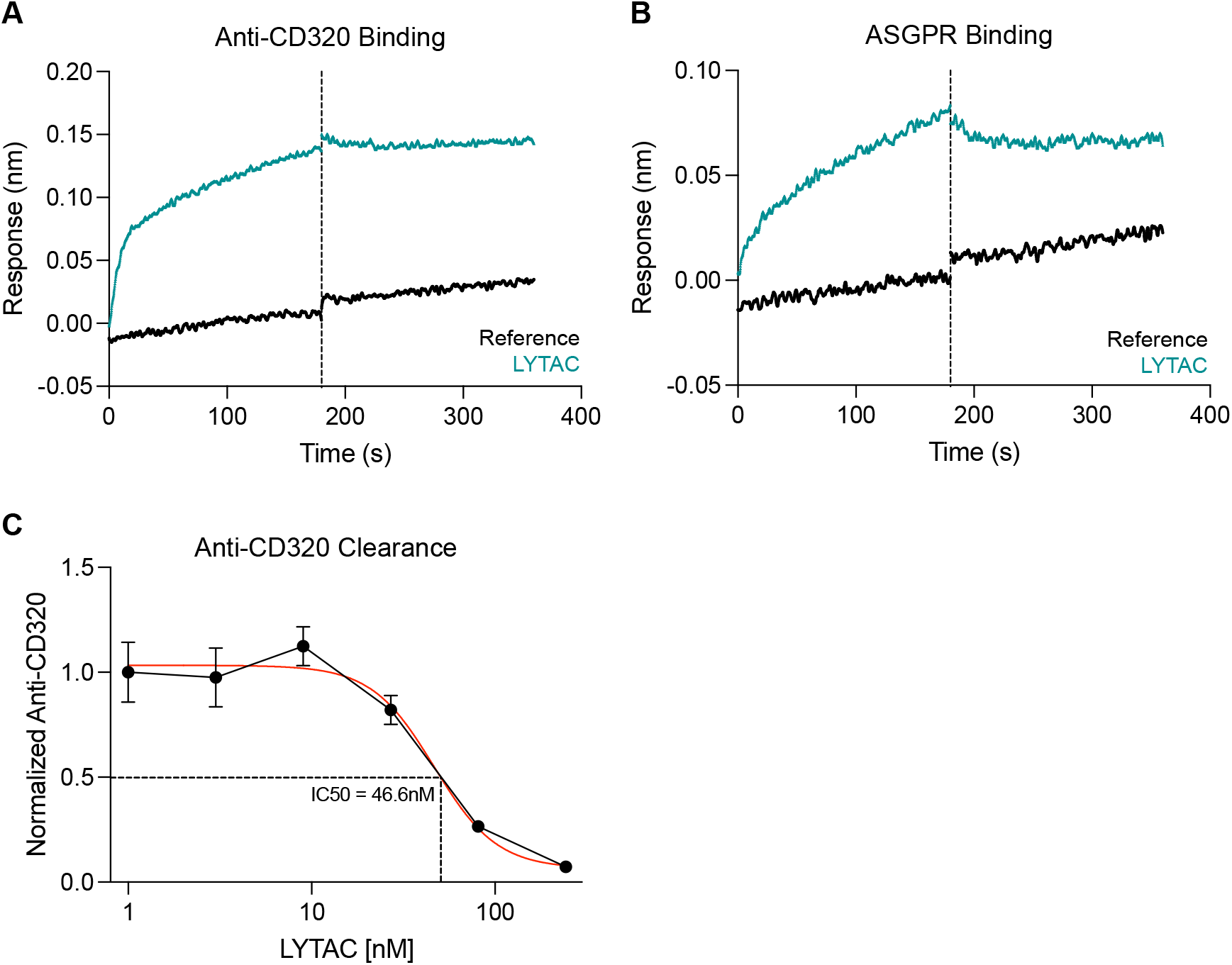
(**A**) Association-dissociation curve of LYTAC + Anti-CD320 interaction measured by biolayer interferometry. (**B**) Association-dissociation curve of LYTAC + ASGPR-Fc interaction measured by biolayer interferometry. (**C**) Dose-response curve showing anti-CD320 concentration in HepG2 supernatant after 24 hours of treatment with LYTAC at increasing concentrations.

